# Circadian timing and entrainment properties of the SCN pacemaker in the PS19 mouse model of tau pathology

**DOI:** 10.1101/2025.06.06.655835

**Authors:** Nicklaus R. Halloy, Megan Formanowicz, Nguyen Nhi Lien Pham, Kari R. Hoyt, Karl Obrietan

## Abstract

Tauopathies are a group of neurodegenerative disorders caused by the misfolded microtubule-associated protein tau (MAPT), leading to its abnormal accumulation and hyperphosphorylation, and resulting in neuronal dysfunction and death. Tauopathy patients also experience disruptions to circadian rhythms of behavior and sleep. The connection between tau pathology and circadian dysfunction is not well understood, especially regarding the role of the suprachiasmatic nucleus (SCN), the brain’s central circadian pacemaker. Here, we conducted histological and functional analyses of the SCN in the PS19 (Prnp-huMAPT*P301S) mouse model of tauopathy. The SCN of PS19 mice had accumulation of phosphorylated tau as early as 2 months of age, and tau pathology was detected in both major neuronal subpopulations of the SCN: VIPergic (core) and AVPergic (shell) neurons. To assess SCN timing and entrainment properties, daily locomotor activity was monitored in PS19 and wild-type (WT) mice from 3 to 11 months-of-age. Activity profiles, rates of re-entrainment to changes in the light/dark cycle, and intrinsic circadian timing properties were unchanged in PS19 mice compared to age-matched WT mice. Finally, profiling circadian gene expression in tau fibril-seeded SCN explants from PS19 and WT mice did not reveal differences in network-level oscillator properties. Together, these findings suggest that tau pathology within the SCN is not sufficient to trigger marked disruptions of core circadian timing mechanisms in this tauopathy model. Further, these results raise the possibility that circadian disruptions in tauopathies arise from dysfunction in SCN-gated output pathways or downstream clock-gated circuits rather than the SCN oscillator itself.

## 1. Introduction

Tauopathies are a class of neurodegenerative disorders characterized by the pathological accumulation of hyperphosphorylated tau protein in the brain. These disorders include Alzheimer’s disease (AD), frontotemporal dementia (FTD), progressive supranuclear palsy, and corticobasal degeneration, all of which manifest varying degrees of cognitive and behavioral impairment (Creekmore et al., 2024; Orr et al., 2017). Further, patients with tauopathies also develop circadian disruptions, such as fragmented sleep (Holth et al., 2016; Sani et al., 2019; Vitiello et al. 1990;) and irregular activity-rest cycles (Anderson et al., 2009; Harper et al., 2005; Hastings and Goedert, 2013; Musiek et al., 2018; Witting et al. 1990), which often worsen as the disease progresses (Musiek, 2017; Musiek and Holtzman, 2016). However, the underlying mechanisms linking tau pathology to circadian dysfunction remain poorly understood, particularly regarding the role of the suprachiasmatic nucleus (SCN), the central pacemaker of the mammalian brain (Hoyt and Obrietan, 2022; Sheehan and Musiek, 2020; Son et al., 2024b).

The suprachiasmatic nucleus (SCN), located in the anterior hypothalamus, serves as the primary circadian pacemaker in mammals. It regulates a wide range of physiological and behavioral rhythms, including locomotor activity, sleep-wake cycles, hormone secretion, and body temperature (Hastings et al., 2018). The SCN exerts its influence in large part by coordinating the phasing and amplitude of peripheral circadian oscillator populations, ensuring highly organized rhythmic output across all organ systems. Circadian rhythms are generated by transcriptional-translational feedback loops involving core clock genes, including *Clock, Bmal1, Per1/2,* and *Cry1/2* (Hastings et al., 2019; Welsh et al., 2010). Disruption of these rhythms is increasingly recognized as a hallmark of neurodegenerative diseases, often correlating with disease progression and severity (Colwell, 2021; Leng et al., 2019). In line with this idea, post-mortem histological analysis identified high levels of phosphorylated tau (p-tau) in SCN neurons of individuals with Alzheimer’s disease (Son et al., 2024a; Swaab et al., 1992). Additionally, a long free-running rhythm phenotype and disrupted expression of Period2 in the SCN were reported in the Tg4510 mouse model of tauopathy (Stevanovic et al., 2017). As such, these findings provide a rationale for additional mechanistic studies that examine the effects of pathological tau on entrainment and timing properties of the SCN. To investigate the impact of tauopathy on circadian function, we profiled clock timing in the PS19 transgenic mouse model, which overexpresses the human P301S mutant form of tau. This model recapitulates key features of human tauopathies, including the accumulation of hyperphosphorylated tau and the formation of neurofibrillary tangles (Yoshiyama et al., 2007). Further, age-dependent pathological progression in PS19 mice is well-characterized: by 3 months of age, PS19 mice exhibit gliosis and synaptic loss; by 6 months, neurofibrillary tangles appear; and by 9 months, significant neuronal loss is observed.

Here, we report tau pathology within the SCN in the PS19 mouse model; notably, phospho-tau levels were detected at a very early stage of the pathogenic process (2 months-of-age). Interestingly, however, the inherent clock timing properties of the SCN are largely intact in the PS19 mouse line-even at the later stages of tau pathology. These, and other data provided here, indicate that the clock timing properties of the SCN are largely resilient to the pathogenic processes driven by tau. Understanding how tau pathology affects clock-gated physiology (e.g., central vs peripheral clocks and which brain regions) is crucial, as circadian disruption might not only a symptom, but could also contribute to disease progression.

## 2. Methods

### 2.1 Transgenic Mouse Models

For wheel-running and immunohistochemistry/immunofluorescence experiments, PS19 breeder pairs (B6;C3-Tg(Prnp-MAPT*P301S) PS19Vle/J Strain #008169; Yoshiyama et al., 2007) were obtained from The Jackson Laboratory (Bar Harbor, ME). Genotyping of PS19 transgenic and WT littermate offspring was performed by Transnetyx (Cordova, TN). Mice were maintained on a 12-h light/dark cycle (lights on at 6:00 a.m., lights off at 6:00 p.m.) with *ad libitum* access to food and water. All animal care and experiments described here were conducted in accordance with the guidelines for animal experimentation at The Ohio State University and were approved by the Institutional Animal Care and Use Committee (IACUC).

For slice culture experiments, female PS19 mice were crossed to male *Per1*-Venus mice (clock fluorescent reporter; C57BL/6 background) to generate PS19 and WT littermates expressing the *Per1*-Venus reporter for real-time circadian profiling of clock rhythms in the SCN (Cheng et al., 2009; Hoyt et al., 2023). Genotyping was performed by Transnetyx (Cordova, TN).

### 2.2 Activity-based Assessment of Circadian Clock Timing and Entrainment

Wheel running activity was used as a measure of SCN clock timing and clock entrainment. At three months of age,16 male mice (8 PS19, 8 WT littermates) were individually housed in clear polycarbonate cages equipped with running wheels (15 cm diameter), magnetic sensors, and *ad libitum* access to food and water. Wheel rotation data was binned into 5-min intervals and collected using the VitalView6 hardware and software system (Starr Life Sciences; Oakmont, PA). Sex differences in physical and behavioral outcomes have been observed in PS19 mice accompanied by increased tau phosphorylation in male PS19 mice (Sun et al., 2020). As such, to assess potential tau pathology-associated sex-related differences in circadian behavior, 12 female mice (6 PS19, 6 WT) were also profiled (beginning at 7 months-of-age) using the noted running wheel assay.

### 2.3 Assessment of Circadian-Regulated Locomotor Activity

Mice were initially entrained and maintained on a 12-h light (300 lux)/12-h dark (LD) cycle (6:00 am lights on, 6:00 pm lights off) for at least 7 days prior to experimental data collection. For profiling of circadian locomotor activity in LD, mice were maintained in this condition for an additional 10 days. To examine the rate of re-entrainment to a shift in the LD cycle, mice were subjected to a 6-h advance and 6-h delay in the phase of the LD cycle. Mice were kept in each of these conditions for at least 10 days to allow adequate time for re-entrainment to the new LD cycle. To assess inherent pacemaker activity under constant conditions, we used 24-h dark (constant darkness: DD) and 24-h light (constant light: LL-300 Lux) paradigms. Mice were maintained under DD for at least 17 days and under LL for at least 14 days.

### 2.4 Evaluation of Circadian Clock Timing and Entrainment

Activity-based assessments were performed with ClockLab Analysis software (ActiMetrics; Lafayette, IN) and Actiview software (Starr Life Sciences; Oakmont, PA). “Overall activity” in LD, DD, and LL was determined by averaging the daily activity counts for each 5-minute binned period for a 9-day period, and the averaged values were calculated as either the total mean activity count for a 24-h period or as a mean 24-h longitudinal activity profile (see Fig. 4B). “Phase angle” indicates the difference in time between the onset of clock-gated locomotor activity and the start of the dark phase of the LD cycle. “Days to entrain” and entrainment profiles during both “phase advance” and “phase delay” were calculated based on the daily, circadian clock-actuated activity onset. Activity onset was defined as the first period in which mice demonstrated locomotor activity for at least 30 consecutive minutes (six-5 min binned periods) that constituted the initiation of the daily bout of consolidated locomotor activity. Entrainment was defined as an activity onset within 15 minutes of the new dark phase with activity onset stabilizing for the next three days. Circadian periodicity (i.e., τ) in DD and LL was determined using the Chi Square Periodogram measurement tool in ClockLab and confirmed using a linear regression method (Actiview). Alpha (activity period duration) was defined as the duration of the clock-gated, consolidated activity period; termination of this period was defined by a break in activity that persisted for > 60 minutes. Acrophase (peak in the activity rhythm) in DD was calculated using ClockLab; the daily activity onset at ∼ CT12 was used as a starting point for the analysis. For Fig 6E, the normalized activity level in LL, and in the LD period following LL (LD_post_), was determined by calculating the percent activity of each mouse under these conditions to the LD_pre_ activity level, which was set = to 100%. Due to the physical decline (e.g., hindlimb weakness) observed in some aged PS19 mice (Yoshiyama et al., 2007) animals were evaluated weekly for overall health. Mice were removed from actimetry experiments and euthanized if paraparesis markedly reduced activity, as assessed in VitalView; in total, 4 PS19 mice; 3 males, 1 female were removed from the study.

All actimetry data were analyzed using Student’s t-test, unless otherwise noted. GraphPad Prism software (Version 10.2.2; San Diego, CA) was used for analysis, and p < 0.05 was defined as a statistically significant difference. In all graphs that include error bars, the data points represent the mean ± <u>S</u>tandard <u>E</u>rror of the <u>M</u>ean (S.E.M.).

### 2.5 Brain Tissue Isolation and Immunolabeling

Mice were euthanized via cervical dislocation followed by rapid decapitation. Brain tissue was isolated, placed into ice cold oxygenated media (120 mM NaCl, 3.5 mM KCl, 2.5 mM CaCl_2_, 1.25 mM NaH_2_PO_4_, 26 mM NaHCO_3_, 1.3 mM MgCl_2_, 10 mM dextrose in deionized water, pH = 7.4) and was then cut into 600-μm-thick coronal slices with a Leica vibratome (catalog #: VT1200, Leica Biosystems, Nussloch, Germany). Sections were transferred to a 4% formaldehyde (F8775, Sigma) solution diluted in phosphate buffered saline (PBS; 21600-069, Gibco) for 6-h at room temperature, then cryoprotected at 4°C overnight in 30% sucrose in PBS (w/v, with 0.02% sodium azide). Sections were thin cut (40 μm) with a sliding microtome (SM2000R, Leica Biosystems) and transferred into FD Section Storage Solution (PC101, FD NeuroTechnologies) for storage at −20°C.

For immunohistochemical (IHC) labeling, SCN-containing sections were blocked and permeabilized in 5% bovine serum albumin (BSA)/PBS-T (PBS containing 0.1% Triton-X100) with 0.02% sodium azide for 90 minutes followed by treatment with 0.3% H_2_O_2_ in PBS for 15 minutes at room temperature. Sections were incubated overnight at 4°C with biotinylated AT8 (mouse host, 1:1000 dilution, catalog #: MN1020B, Invitrogen, Waltham, MA) antibody in 5% BSA/PBS-T with 0.02% sodium azide. Tissue was then processed with a VECTASTAIN ABC-HRP kit (PK-4000, Vector Laboratories, Newark, CA) per the manufacturer’s instructions, and the signal was visualized using nickel-intensified diaminobenzidine (DAB) substrate (SK-4100, Vector Laboratories). Tissue sections labeling for AT8 were developed for 10 min in DAB substrate. Development was terminated by washing the sections 3 times in deionized (DI) water, and sections were then mounted on slides, coverslipped, and sealed with Permount media (SP15, Fisher Scientific). Between each labeling step prior to DAB development, sections were washed in PBS-T (3 washes, 5 min/wash). Brightfield images were captured using a Leica DFC450C camera (Leica Microsystems, Wetzlar, Germany) connected to a Leica DMIRB microscope using 4x objective for whole-section images and 10x objective for regional images (e.g., suprachiasmatic nucleus, piriform cortex, striatum, and retrosplenial cortex). Metamorph imaging software (Version 7.10.2.240; Molecular Devices, San Jose, CA) was used for image acquisition.

To quantify the intensity of AT8 immunohistochemical labeling, a densitometric approach was employed using Fiji/ImageJ (Schindelin et al., 2012). To this end, all images (2560 x 1920 pixels) were converted from a 16- to 8-bit scale (0-255 units), light-dark inverted, and then a cutoff threshold (from 80-255) was applied to define p-tau positive cells. Next, a digital circle (i.e., region-of-interest) was placed in the central SCN (∼ 450 × 450 pixels), and the integrated value of all pixels above the threshold was calculated. For comparisons of p-tau expression in the SCN as a function of age, a one-way ANOVA with Tukey post-hoc test was performed.

For immunofluorescence (IF) labeling of SCN-containing coronal brain sections, tissue was blocked and permeabilized in 3% BSA/PBS containing Triton-X100 (0.2%) and sodium azide (0.02%). Sections were incubated (overnight, 4°C) with 3% BSA/PBS (with 0.2% Triton-X100 and 0.02% sodium azide) containing biotinylated AT8 (mouse host, 1:1000 dilution, catalog #: MN1020B, Invitrogen), AVP (rabbit host, 1:1250 dilution, AB1565, EMP Millipore, Darmstadt, Germany), and/or VIP (rabbit host, 1:250 dilution, RP1108, Boster Biological, Pleasanton, CA) antibodies. Sections were then incubated for 4 hours at room temperature with Alexa 594 conjugated streptavidin (1:1000 dilution, S32356, Invitrogen) and anti-rabbit Alexa 488 (1:1000 dilution, A11034, Invitrogen) in PBS. Between each labeling step, sections were washed in PBS (3 washes, 5 min/wash), unless otherwise noted. Sections were mounted on slides with Fluoromount-G (0100-01, Southern Biotech, Birmingham, AL), and confocal fluorescence images were acquired with a Leica SP8 confocal system (Leica, Nussloch, Germany) using Alexa 488 and Alexa 594 pre-set acquisition settings.

### 2.5 SCN Explant Culture and Fluorescence Profiling of Clock Rhythms

SCN tissue isolated from postnatal day 1 pups from *Per1*-Venus mice crossed with PS19 mice was used for imaging-based timing assays. Brain tissue was isolated, and 200-μm-thick coronal sections through the SCN and surrounding hypothalamic tissue were prepared using a McIlwain Tissue Chopper. SCN-containing sections were isolated, and the SCN was resected with microdissection scissors. SCN tissue was washed 2 times in Hibernate-A medium (A1247501, Gibco) and then incubated for 30 min with 2 mg/ml (∼50 units) of papain suspension (LS003126, Worthington Biochemical) at 35°C. After proteolytic digestion, tissue was washed 2 times with Hibernate-A, then triturated into large (50-200 um) intact pieces of SCN tissue in Neurobasal-A media (1088022, Gibco) containing 2% B27-plus supplement (with antioxidants; A3582801, Gibco), 100 U/ml penicillin/100 μg/ml streptomycin (5140122, Gibco), and 0.5 mM GlutaMAX-1 (35050061, Gibco). At this point, explant SCN tissue was plated in 48-well culture plates (83.3923.300, Sarstedt) coated with poly-D-lysine (0.1 mg/ml for 1 hour; A38904, Gibco). Cultures were maintained at 35°C and 5% CO_2_ in a cell culture incubator until long-term rhythms profiling began (2-9 days after culturing). Genotyping of each pup was performed by Transnetyx (Cordova, TN).

At the beginning of the long-term imaging assays, culture plates were sealed with an ALA MEA-SHEET (Multi Channel Systems, Reutlingen, Germany) then moved to a stage-top incubator (Oko Labs, Pozzuoli, Italy), set to 5% CO_2_, 35°C and 95% humidity, located on an inverted Leica Stellaris confocal microscope (Leica, Nussloch, Germany). Long-term imaging of Venus expression was performed using YFP (yellow fluorescent protein) acquisition settings, and time-lapse based image acquisition was performed at 1-h intervals for up to 6 weeks (for a detailed description of the imaging methods, see Wheaton et al., 2018 and Hoyt et al., 2023). For tau seeding experiments, after baseline acquisition, SCN explants were treated with 1.7 µg/ml or 7.0 µg/ml human recombinant Tau-441 (2N4R) P301S mutant protein pre-formed fibrils (tau PFF; Stressmarq SPR-329).

*Per1*-Venus expression profiling of SCN explants was performed using Fiji/ImageJ (Schindelin et al., 2012), and Biodare2 (Zielinski et al., 2014; Zielinski et al., 2022) was used to linear detrend the Venus time-lapse data and calculate the period of cellular Venus expression (FFT-NLLS method) during acquisition. One-way ANOVA was used to compare baseline period to tau PFF-treated periods for each tau PFF concentration for PS19 or WT SCN explants.

To assess tau pathology in the tau PFF-treated SCN explants, the explants were fixed with 4% formaldehyde/0.1% Triton-X100 (in PBS) for 30 minutes then washed 2 times with PBS at the completion of the Venus fluorescence time-lapse experiments. Next, the explants were blocked and permeabilized for 1-h in 3% BSA/0.1% Triton X-100 and 0.2% sodium azide and then incubated with the following primary antibodies: biotinylated AT8 (mouse host, 1:500 dilution, catalog #: MN1020B, Invitrogen), MC1 (mouse host, 1:750 dilution, originally generated by Dr. Peter Davies’ research group; Jicha et al., 1997), and GFP (chicken host, 1:1500 dilution, 13970, Abcam) for 48 hours at 4°C in the same blocking buffer. Explants were then incubated with Alexa 594-conjugated streptavidin (1:500 dilution, S32366, Invitrogen), Alexa 405-conjugated anti-mouse (1:500 dilution, A31553, Invitrogen), and Alexa 488-conjugated anti-chicken (1:500 dilution, A11039, Invitrogen) secondary antibodies in PBS for 48-h. Explants were then incubated with the DNA label DRAQ5 (1:5000 dilution, 62254, Thermo Scientific) for 1 hour. Between each labeling step, sections were washed in PBS (6 washes: three 10 min washes, followed by three 60 min washes). Fluorescence images of the immunolabelled cells were acquired with an inverted Leica Stellaris confocal microscope (Leica, Nussloch, Germany) using Alexa 405 (MC1), Alexa 488 (GFP for Venus), Alexa 594 (AT8), and DRAQ5 (nuclei) pre-set acquisition settings and a 10x objective.

## 3. Results

### 3.1. The PS19 Transgenic Mouse Line

For the *in vivo* examination of the relationship between tauopathy and the timing properties of the SCN, we used the PS19 transgenic mouse model. In this model, a P301S mutation (proline to serine at codon 301) within the human microtubule-associated protein tau (MAPT) gene (1N4R isoform) is expressed under the control of the mouse prion protein promoter. This construct leads to high levels of mutant human MAPT expression in the central nervous system (Holmes et al., 2014; Yoshiyama et al., 2007). Notably, these mice exhibit many of the key pathological features associated with Alzheimer’s disease and familial frontal temporal lobe dementia. Along these lines, PS19 mice develop age-dependent tau hyperphosphorylation, aggregation, and formation of neurofibrillary tangles associated with neuronal loss, reactive astrogliosis, and behavioral deficits (Patel et al., 2022; Takeuchi et al., 2011; Yoshiyama et al., 2007). PS19 transgenic mice exhibit weak expression of perikaryal and dendritic tau within forebrain regions, including the hippocampus and amygdala, as well as in the brain stem and spinal cord (Yoshiyama et al., 2007). Consistent with this, the first pathological effects of the PS19 transgene include synaptic loss and reactive gliosis at ∼ 3 months of age with neurofibrillary tangles and cognitive impairment arising at ∼ 6 months of age.

### 3.2. Phosphorylated Tau Expression in the SCN of PS19 Mice

To examine transgenic tau expression, we used the AT8 antibody, which labels human tau protein phosphorylated (p-tau) at Serine 202 (Ser202) and Threonine 205 (Thr205; Goedert et al., 1995; Malia et al., 2016). Phosphorylation of tau at Ser202/Thr205 is associated with disrupted microtubule binding of tau proteins, aggregation of dissociated tau, and the development of paired helical filaments, which are the predominant component of neurofibrillary tangles (Despres et al., 2017; Kidd, 1963; Mammeri et al., 2024; Moloney et al., 2021). Coronal brain sections from PS19 mice labelled with the AT8 antibody (Fig. 1A-1C) revealed a progressive age-dependent accumulation of p-tau labeling, which was most notable within cortical structures. Interestingly, within the SCN, diffuse, yet marked, p-tau expression was observed from 2 months to 11 months-of-age (Fig. 1A’-1C’; control p-tau labeling in WT aged-matched SCN sections are shown in panels E-G). Of note, 2 months-of-age is a developmental time point that precedes the reported onset of pathological alterations in CNS physiology (Yoshiyama et al., 2007). Further, with increased age (Fig. 1B’ and 1C’), a subset of SCN cells exhibited very high levels of AT8 labeling, which is consistent with the formation of tau-positive inclusions (Hamlin et al., 2024; Moloney et al., 2021). Quantitative analysis of SCN labeling (Fig 1H) shows similar levels of p-tau at 2 and 4 months, with a decrease in overall labeling at 11 months, possibly reflecting an age— related changes in the phosphorylation patterns of pathogenic tau (Hamlin et al., 2024; Rawat et al., 2022).

**Figure 1.**
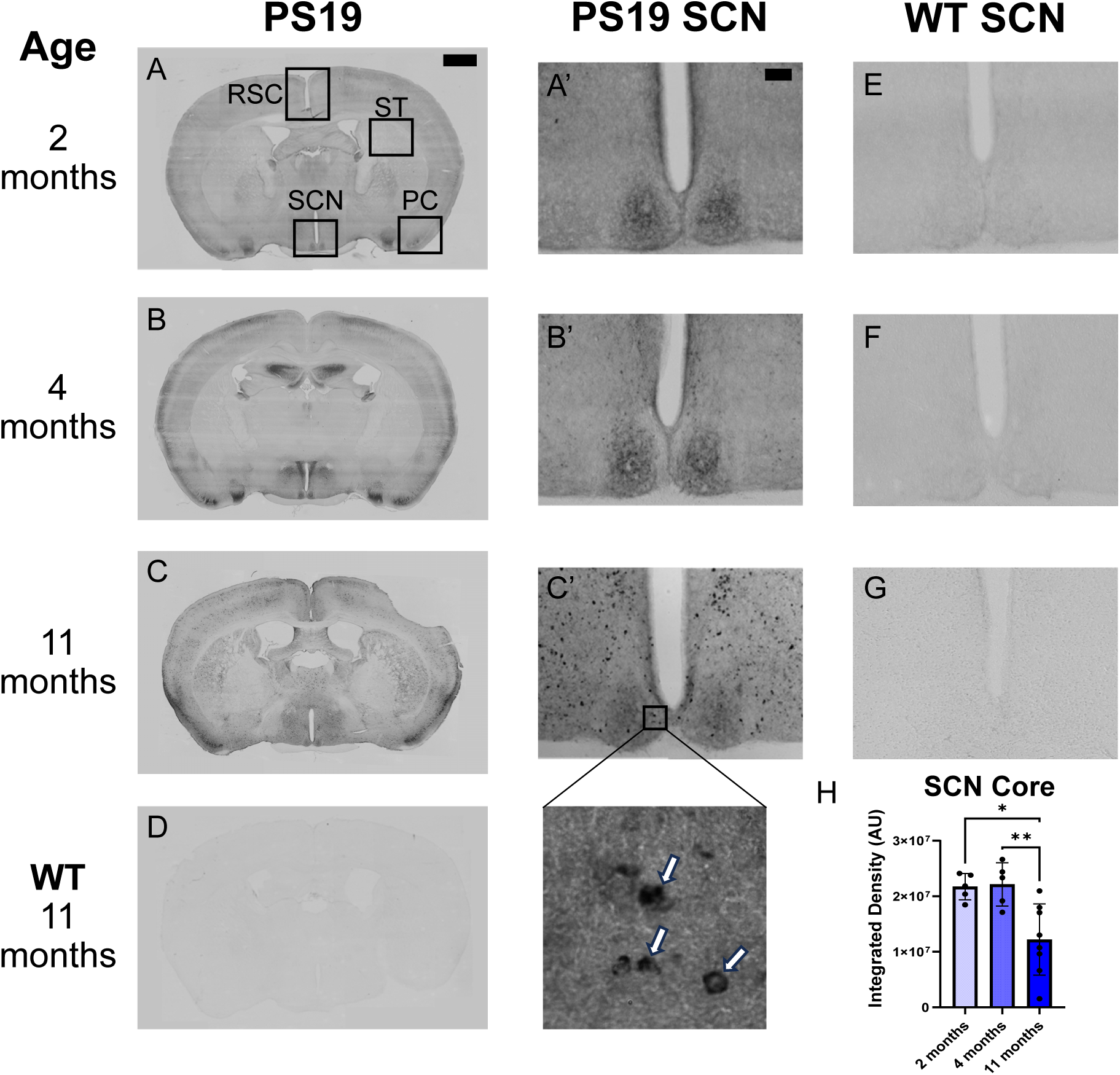
Phosphorylated tau (p-tau) is elevated in the suprachiasmatic nuclei (SCN) of PS19 mice. Representative coronal brain sections from 2-, 4-, and 11-month-old PS19 (A-C’) and an 11-month-old WT littermate (D) labeled with the AT8 antibody recognizing p-tau and developed for immunohistochemistry using a DAB-based method. Dark labeling represents AT8 antibody labeling (Boxed regions in A: SCN: suprachiasmatic nuclei, RSC: retrosplenial cortex, ST: striatum, PC: piriform cortex). Higher magnification images of the SCN reveal marked p-tau labeling at 2-months of age (A’) with small inclusions beginning to form in the SCN by 4 months (B’), and by 11 months (C’) inclusions are broadly observed within the SCN and the surrounding hypothalamic tissue (arrows in magnified image denote cells with inclusions). E-G: Age-matched WT littermate brain tissue was immunolabeled using the AT8 antibody for comparison, and very low DAB reactivity was detectable, as expected. H: Quantification of the AT8 signal in the SCN of PS19 mice at 2-, 4-, and 11-months with labeling at 11 months significantly lower than 2- and 4-month-old SCN, * = p < 0.05, ** = p < 0.01, ANOVA; 5-8 animals per group. Scale bars in panels A and A’ are 1000 and 100 microns, respectively.

To advance our analysis, p-tau accumulation was examined within the piriform cortex, striatum and retrosplenial cortex (Fig. 2). Consistent with prior studies (Ahmad et al., 2021; Chen et al., 2025; Wagner et al., 2015), a marked accumulation of p-tau was observed in all the noted brain regions and followed a distinct region-specific temporal profile. Along these lines, low levels of expression were detected in the piriform cortex at 2 months-of-age, and by 11 months-of-age, there was a marked increase in both the number of p-tau positive cells and the intensity of labeling (Fig 2A-2C). Interestingly, within the striatum and retrosplenial cortex, limited p-tau expression was detected at 2 and 4 months of age, whereas at 11 months, marked accumulation could be observed in both brain regions (Figure 2D-2I).

**Figure 2.**
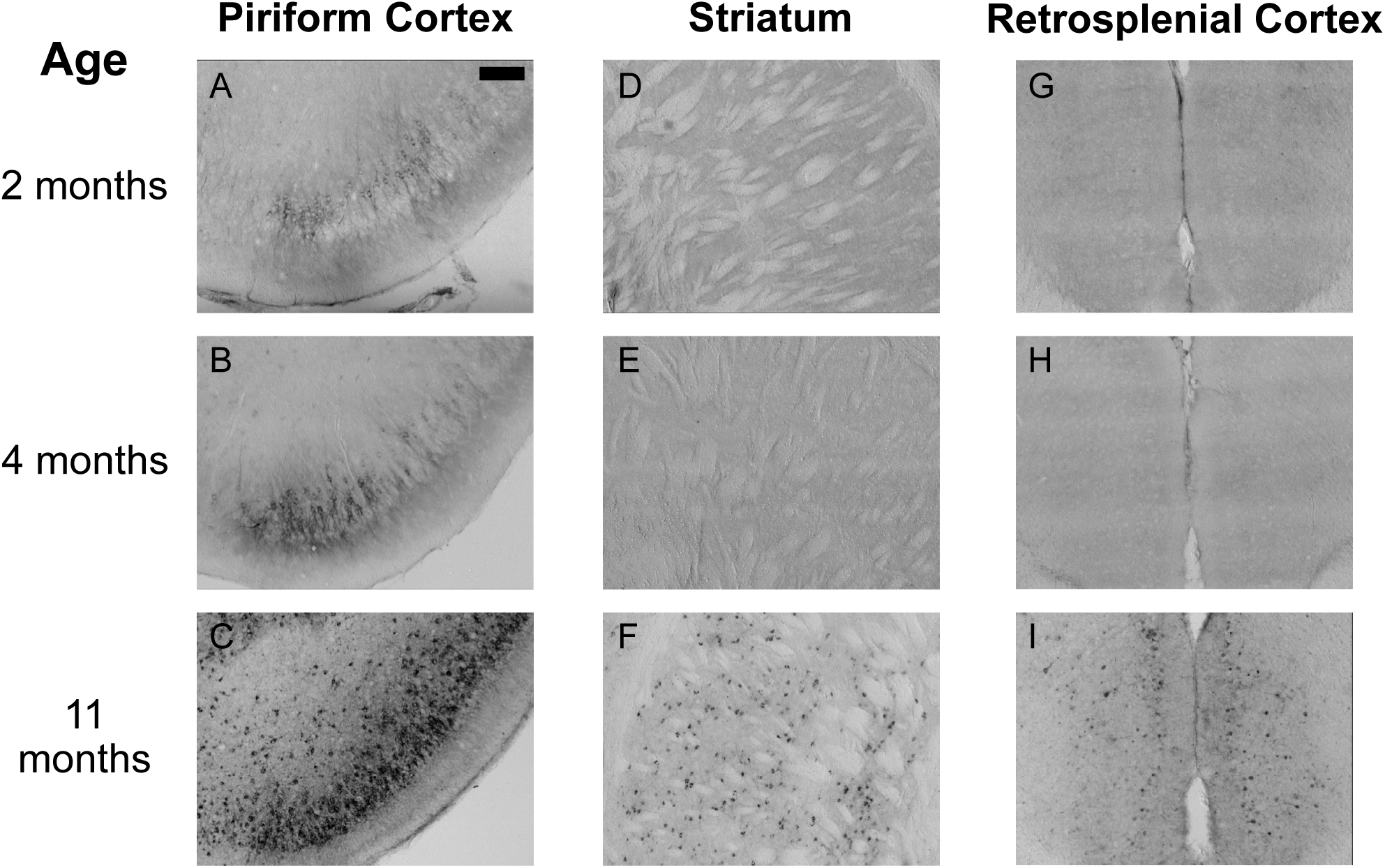
Cortical and striatal expression of p-tau increases with age in PS19 mice. Immunohistochemical labeling for p-tau (from Fig 1; AT8 antibody) in the piriform cortex (A-C), striatum (D-F), and retrosplenial cortex (G-I) in 2-, 4-, and 11-month-old PS19 mice. P-tau is observed in the piriform cortex at 2 months-of-age (A) with a marked increase in labeling at 4 months-of-age (B), and inclusions are detected at the 11-month time point (C). The striatum and retrosplenial cortex have clear p-tau labeling at 11 months-of-age, but not at 2- or 4-months-of-age (D-I). Scale bar in panel A is 200 microns.

Next, fluorescence-based immunolabeling was used to profile AT8 labeling within major subpopulations of SCN neurons. To this end, tissue from 9-month-old WT and PS19 mice was co-labeled for AVP (arginine vasopressin), a marker of neurons within the dorsal and medial regions of the SCN (referred to as the SCN shell; Fig. 3A-F’), and VIP (vasoactive intestinal peptide), a marker of ventral and lateral regions of the SCN (referred to as the SCN core; Fig. 3G-L’). Coronal sections of SCN from PS19 mice revealed the expected expression pattern and levels of AVP and VIP within the SCN. As anticipated (based on patterns of DAB immunolabeling, Fig. 1), p-tau was colocalized with both AVP- and VIP-positive neuronal cell populations within the SCN.

**Figure 3.**
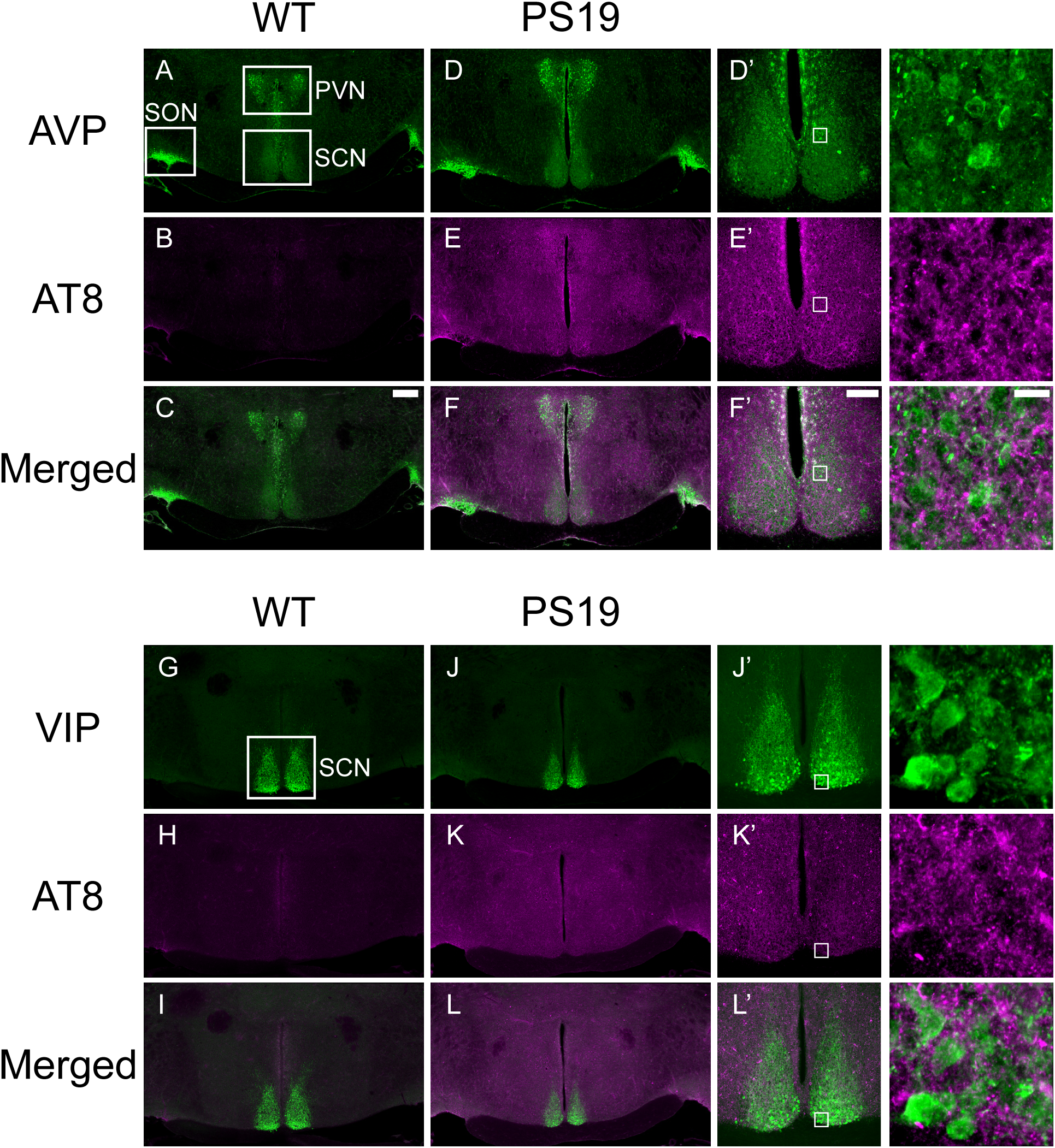
Tau pathology in SCN shell and core neuronal populations in PS19 mice. Immunofluorescence labeling for p-tau (AT8 antibody; magenta) and arginine vasopressin (AVP; a marker of SCN shell neurons; A-F’; green) or vasoactive intestinal peptide (VIP; a marker of SCN core neurons; G-L’; green) from 9-month-old WT littermate and PS19 transgenic mice. Confocal images were generated from 40-μm- thick sections of the central SCN. Boxed regions in D’-F’ and J’-L’ are magnified and shown in the panels to the right. The magnified images were generated from maximum-projected z-stacks of the 40-μm thick section and are provided to highlight the colocalization of phosphorylated tau in AVP- and VIP-positive neurons. SCN: suprachiasmatic nucleus; SON: supraoptic nucleus; PVN: paraventricular nucleus. Scale bar in panel F’ is 200 microns; and 10 microns in the magnified panel to the right.

Together, these data reveal a marked early-onset expression pattern of pathological transgenic tau within the SCN. This, in some respects, is similar to recent findings from humans showing early-stage SCN tau pathology in AD patients (Son et al., 2024a). As such, these data provide a strong rationale for examining the SCN-based circadian timing properties of PS19 mice.

### 3.3. SCN Clock Timing and Entrainment in PS19 Mice

Next, we examined the SCN clock timing properties of the PS19 mouse line. To this end, we used locomotor activity (wheel running) as a readout of circadian periodicity and phasing under a number of entrainment and free-running paradigms. Of note, sex-related differences in pathogenesis severity and functional outcomes have been observed in humans and murine models of tauopathy (Oveisgharan et al., 2018; Sun et al., 2020). Apart from a modest, yet significant, difference in acrophase and alpha in male vs female WT mice (9-10 months old), no other sex-related differences were detected in any measured circadian parameters (Supplemental Table 1). Therefore, data from male and female mice were pooled for group comparisons based on age and genotype.

Initially, we profiled activity under a standard 12-h light:dark cycle (12:12 LD). Representative double plotted actographs from PS19 and WT littermates (starting at 3 and 8 months-of-age; Figure 4A) demonstrate the expected activity pattern, with the majority of locomotor activity occurring during the dark phase. The average activity profile, the ratio of night/day activity, and the total amount of activity were not significantly different between the PS19 and WT mice at each age (Fig. 4B-D). Furthermore, activity onset (i.e., phase angle of entrainment) was not significantly different between the PS19 and WT mice at the noted ages (Fig. 4C-D). Together, these data suggest that activity levels, photic entrainment capacity (i.e., the strength of the zeitgeber) and the periodicity of the oscillator were not affected by the expression of pathologic tau.

**Figure 4.**
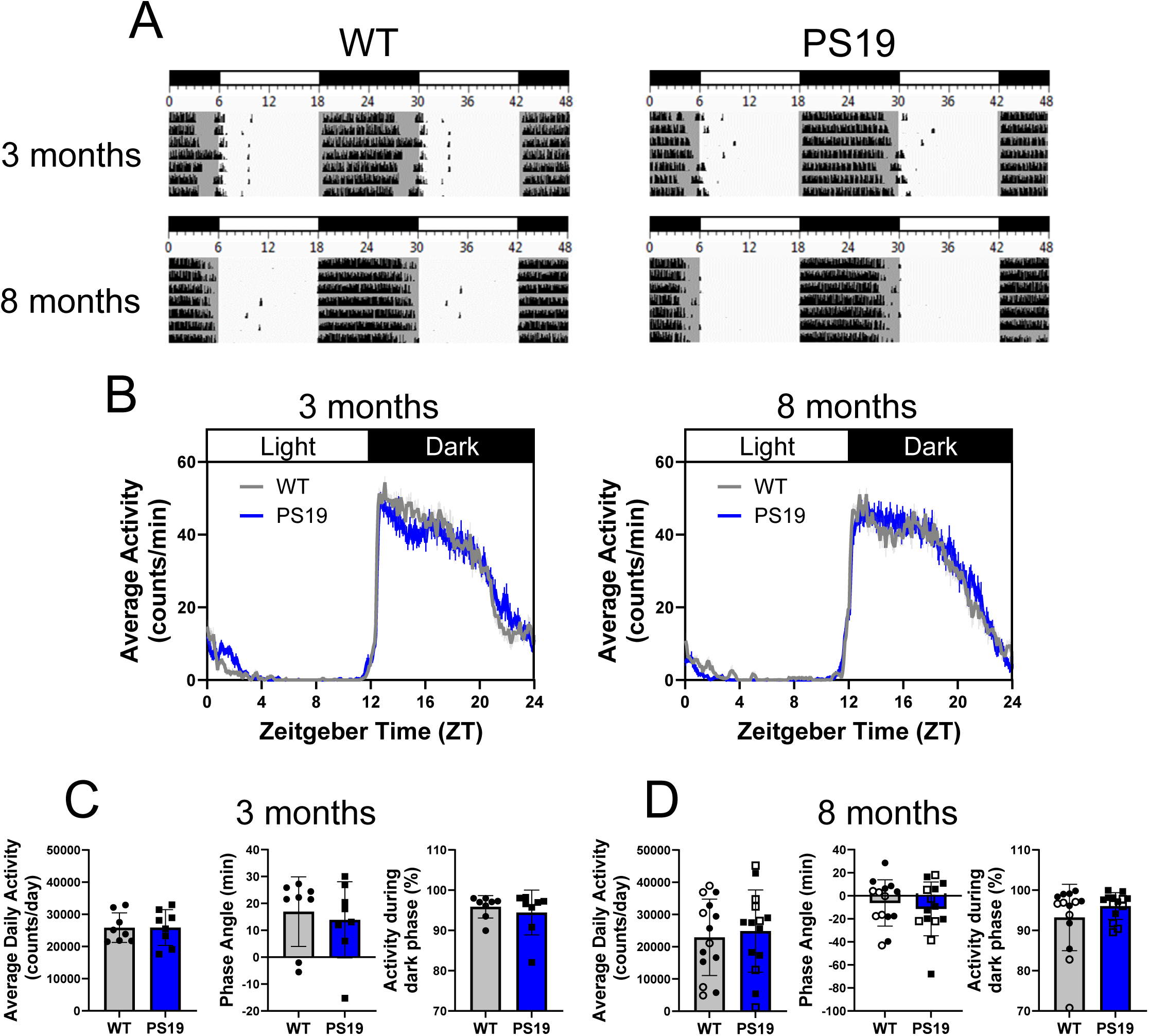
Circadian-gated locomotor activity in PS19 mice. (A) Representative double plotted actograms of wheel running activity from one WT and one PS19 mouse in a 12:12 h light:dark (LD) condition at 3 months- and 8 months-of-age. Darkened regions of the actograms denote periods when the lights were off. (B) Mean (± SEM) daily activity profiles of WT (grey) and PS19 (blue) mice in LD. Quantification of the mean (± SEM) average daily activity (counts/day), phase angle of entrainment of activity onset relative to the start of the dark phase, and percent of total activity in the dark phase for 3-month-old (C) and 8-month-old (D) mice. Data in B-D were analyzed from 8-14 mice for each genotype. Circadian entrainment and locomotor activity were comparable between PS19 mice and WT littermates at both ages tested (no statistically significant changes were identified for the noted parameters at each age; Student’s t-test). Closed shapes indicate male mice (both 3- and 8-month-old) and open shapes indicate female mice (8 month-old).

To further examine clock entrainment, mice were exposed to a 6-h phase resetting paradigm. The 12:12 LD cycle was initially advanced 6-h and then delayed 6-h (Fig. 5A), and the rates of re-entrainment to the advance and delay in the light cycle were quantified (Fig. 5B & C). An examination of the days to re-entrain did not detect significant differences between WT and PS19 mice at 3-4 or 8-9 months-of-age. Further, no significant differences were detected in the daily rate of re-entrainment (based on the mean daily phase entrainment profiles plotted in Fig. 5B & C). Together, these data further support the idea that p-tau accumulation and aggregate formation within the SCN does not affect the resetting capacity of the SCN to light.

**Figure 5.**
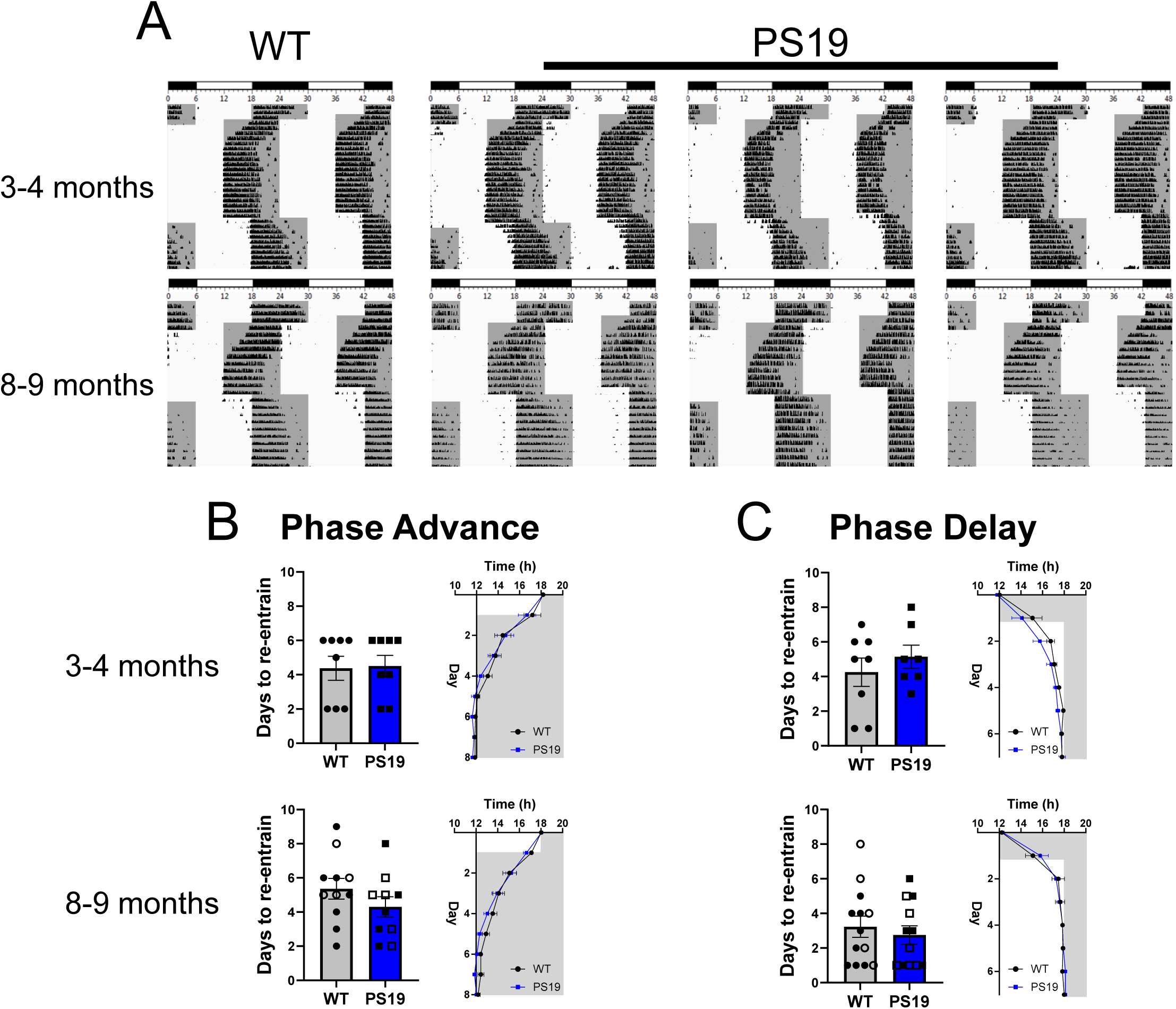
Photic re-entrainment in PS19 mice. (A) Representative double plotted actograms of wheel running activity from one WT mouse and three PS19 mice at 3-4 months- and 8-9 months-of-age. Mice were entrained to a 12:12 L:D cycle and then were exposed to a 6-h phase advance. After stable re-entrainment following phase advance, mice were subjected to a 6-h phase delay. Darkened regions of the actograms denote periods when the lights were off. Relative to WT animals, PS19 mice did not exhibit significant differences in the rate of re-entrainment to either the light advancing (B) or delaying (C) paradigm. Quantification of the total days for mice to re-entrain and the average re-entrainment profiles in mice at 3-4 months- and 8-9 months-of-age. Data in B-C were collected from 8-13 mice for each genotype. No statistically significant changes were identified between the genotypes for the noted parameters at each age; Student’s t-test. Closed shapes indicate male mice (both 3-4- and 8-9-month-old) and open shapes indicate female mice (8-9-month-old).

To assess the inherent time-keeping capacity of the SCN, mice were transferred from the 12:12 LD cycle to both total darkness (DD) and constant light conditions (LL). Under these conditions (i.e., absent any daily entrainment cues), the periodicity of the locomotor activity can be used as a readout of the inherent pacemaker activity of the SCN, with constant dark conditions typically resulting in a period length that is shorter than 24 hours and constant light leading to a lengthening of the period (and a damping of locomotor activity). Representative data in Fig. 6A depict the free running periods of mice (4-5 months- and 9-10 months-of-age) under conditions of total darkness. Notably, no significant difference in period length was detected between the WT and PS19 mice (Fig. 6B & C). Furthermore, other measures of clock activity, including acrophase (peak activity time), alpha (duration of activity), and daily activity were not significantly affected by transgenic expression of mutant human tau in PS19 mice.

**Figure 6.**
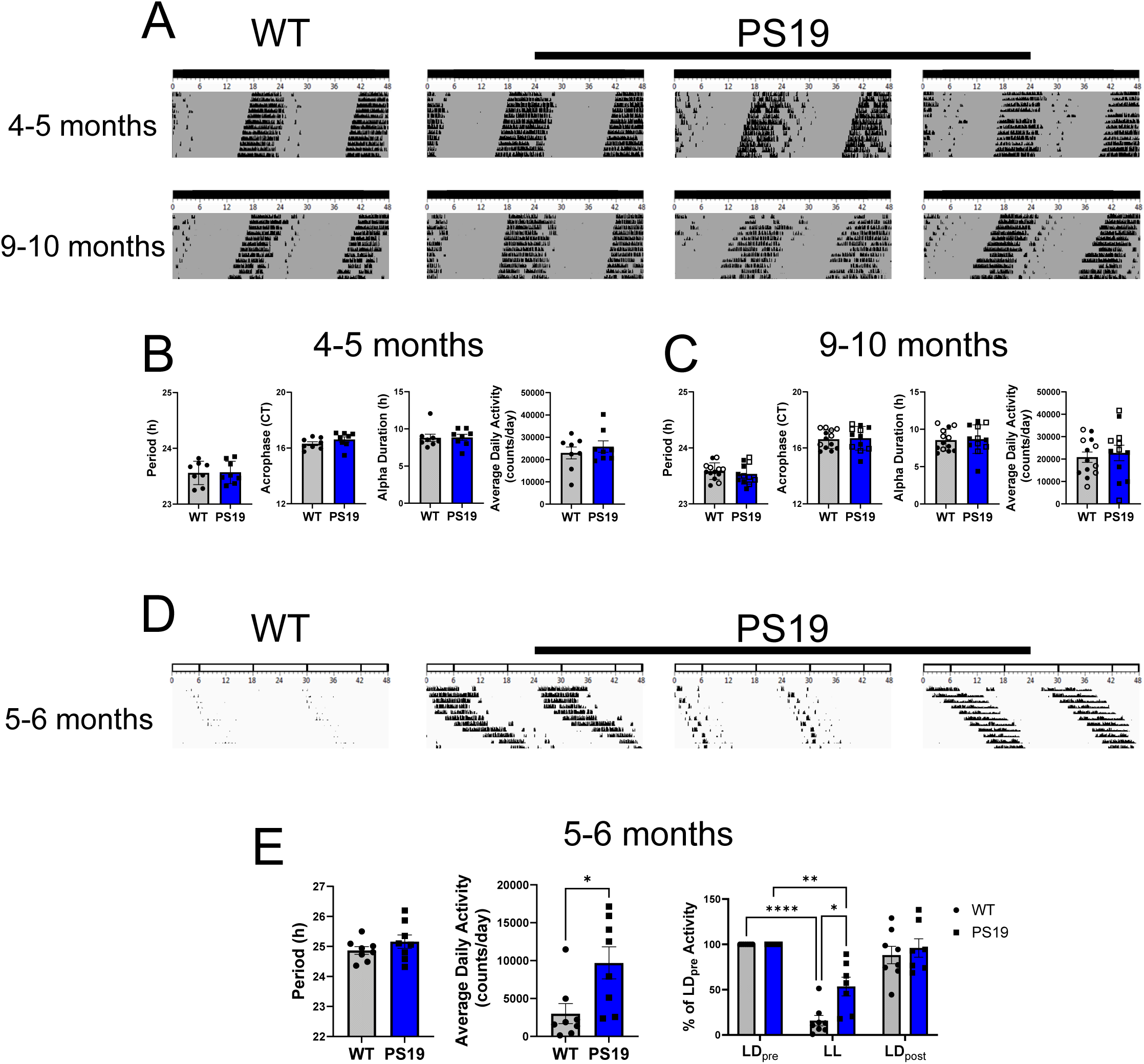
Circadian locomotor activity of PS19 mice under constant dark and light conditions. Representative double plotted actograms of wheel running activity from one WT mouse and three PS19 mice in constant darkness (A) at 4-5 months- and 9-10 months-of-age and constant light (D) at 5-6 months-of-age. (B & C) Quantification of the mean (± SEM) circadian period, acrophase, alpha, and average daily activity in DD at 4- 5 months- (B) and 9-10 months-of-age (C). (E) Quantification of the mean (± SEM) circadian period, average daily activity in LL, and the percentage of activity in LL (relative to activity levels under LD: please see the *Methods* section for a description of the analysis approach). Of note, relative to WT mice, the PS19 mice exhibited a significant reduction in LL-induced activity damping. The percentage activity levels for LL were calculated based on levels of activity in the LD period preceding the transition to the LL condition; The percentage activity is also shown for the LD cycle that followed LL. (E). Data in B & C and E were calculated from 7-13 mice for each genotype; * = p < 0.05, significantly different from WT control; Student’s t-test. Percent of LD_pre_ activity was calculated using 2-way repeated measures ANOVA; * = p < 0.05, ** = p < 0.01, **** = p < 0.0001. Closed shapes indicate male mice (both 3- and 8-month-old) and open shapes indicate female mice (8-month-old).

Under constant light conditions (∼300 Lux), PS19 mice exhibited the expected lengthening of the locomotor activity period and a reduction in overall activity (relative to LD conditions; Daan and Pittendrigh, 1976; Pittendrigh, 1960; Fig. 6D & E). Within age groups, the LL-mediated period lengthening was not significantly different between PS19 and WT mice; however, relative to age-matched WT mice, PS19 mice exhibited a reduced damping of daily activity at 5-6 months (Fig. 6E). Together, the results from the DD and LL assays indicate that inherent clock timing in PS19 mice is not altered as a result of tau hyperphosphorylation and aggregation within the SCN.

A summary of the circadian and entrainment properties of the PS19 and WT mouse lines is presented in Table 1.

**Table 1.** Circadian locomotor activity parameters in early (3-6 months-of-age) and intermediate-late (7-10 months-of-age) stage tau pathology. For parameters 1, 5, and 9, ‘overall activity’ refers to the mean wheel-running activity values per binned 5 min period. Data were averaged over a 9-day period. For parameters 3 and 4, ‘days to re-entrain’ is defined as the number of days needed for animals to be entrained to the new LD cycle. The day when entrainment is completed is defined as the first day when the time between dark onset and activity onset is < 15 min and stabilizes for the next 3 days. For parameter 7 ‘duration of alpha’ refers to the time running bouts persisted without a break of > 60 minutes. Data are presented as the mean ± SEM for all values. * = p < 0.05; statistically significant changes in PS19 mice compared to WT were assessed via Student’s t-test.

| Pathological Stage | Parameter (mean ± standard error) | Wild-type (n) | PS19 (n) | P-Value |
| --- | --- | --- | --- | --- |
| Early | 1. Overall activity in LD (counts/5 min) | 89.70 ± 5.66 (8) | 89.83 ± 6.83 (8) | 0.989 |
|  | 2. LD Phase Angle of activity onset relative to lights off (min) | 16.92 ± 4.59 (8) | 13.89 ± 5.01 (8) | 0.662 |
|  | Positive values indicate activity onset after “lights off” |  |  |  |
|  | 3. Days to re-entrain to 6h advancing LD cycle (d) | 4.38 ± 0.71 (8) | 4.50 ± 0.63 (8) | 0.900 |
|  | 4. Days to re-entrain to 6h delaying LD cycle (d) | 4.25 ± 0.82 (8) | 5.14 ± 0.67 (7) | 0.422 |
|  | 5. Overall activity in DD (counts/5 min) | 79.77 ± 9.25 (8) | 89.65 ± 8.88 (8) | 0.812 |
|  | 6. Circadian period (τ) during DD | 23.56 ± 0.07 (8) | 23.57 ± 0.07 (8) | 0.914 |
|  | 7. Duration of alpha during DD (h) | 8.34 ± 0.19 (8) | 8.86 ± 0.40 (8) | 0.286 |
|  | 8. Acrophase during DD (CT) | 16.31 ± 0.14 (8) | 16.61 ± 0.19 (8) | 0.223 |
|  | 9. Overall activity in LL (counts/5 min) | 10.41 ± 4.58 (8) | 33.68 ± 7.34 (8) | 0.018* |
|  | 10. Circadian period (τ) during LL | 24.77 ± 0.14 (8) | 24.84 ± 0.32 (8) | 0.855 |
| Intermediate-Late |  |  |  |  |
|  | 1. Overall activity in LD (counts/5 min) | 79.57 ± 11.02 (14) | 86.31 ± 11.84 (14) | 0.684 |
|  | 2. LD Phase Angle of activity onset relative to lights off (min) | -6.211 ± 5.36 (14) | -11.38 ± 6.25 (14) | 0.536 |
|  | Negative values indicate activity onset prior to “lights off” |  |  |  |
|  | 3. Days to re-entrain to 6h advancing LD cycle (d) | 6.36 ± 0.61 (11) | 5.3 ± 0.60 (10) | 0.228 |
|  | 4. Days to re-entrain to 6h delaying LD cycle (d) | 3.23 ± 0.61 (13) | 2.75 ± 0.54 (12) | 0.305 |
|  | 5. Overall activity in DD (counts/5 min) | 67.99 ± 8.69 (14) | 79.00 ± 11.93 (12) | 0.455 |
|  | 6. Circadian period (τ) during DD | 23.58 ± 0.04 (13) | 23.54 ± 0.05 (12) | 0.467 |
|  | 7. Duration of alpha during DD (h) | 8.56 ± 0.40 (13) | 8.68 ± 0.55 (12) | 0.857 |
|  | 8. Acrophase during DD (CT) | 16.62 ± 0.18 (13) | 16.69 ± 0.25 (12) | 0.839 |

### 3.4. In vitro SCN Profiling of Clock Rhythms in PS19 Mice

To further assess the effects of tau pathology on SCN timing properties, we turned to an *in vitro* fluorescence-imaging based profiling approach. To this end, the PS19 line was crossed with our transgenic *Per1-*Venus reporter mouse (Cheng et al., 2009; Hoyt et al., 2023), and SCN explants were examined using time-lapse confocal microscopy. Of note, the *Per1*-Venus mouse line drives the expression of a nuclear localized, short-half-life Venus transgene from the *Period1* promotor, thus allowing for a robust and dynamic cellular-level fluorescent readout of the molecular feedback loop that forms the basis of the circadian timing process (Cheng et al., 2009; Hoyt et al., 2023).

SCN explants were cultured from *Per1-*Venus::WT and *Per1-*Venus::PS19 mice and profiled at 1-h intervals over a multiweek period; assessment of circadian period was made in 1-week intervals with the baseline period defined over a 5-day time interval (‘Week 0’) preceding the addition of tau pre-formed fibrils (PFF) for seeding experiments. Representative Venus traces reveal that cultured SCN explants exhibited robust and persistent daily rhythms (Fig. 7O and 7P). Within the “baseline” timeframe, the periodicity of Venus rhythms did not differ between PS19 and WT explants (Fig. 7A-F) indicating that the clock is not affected by transgenic tau expression.

**Figure 7.**
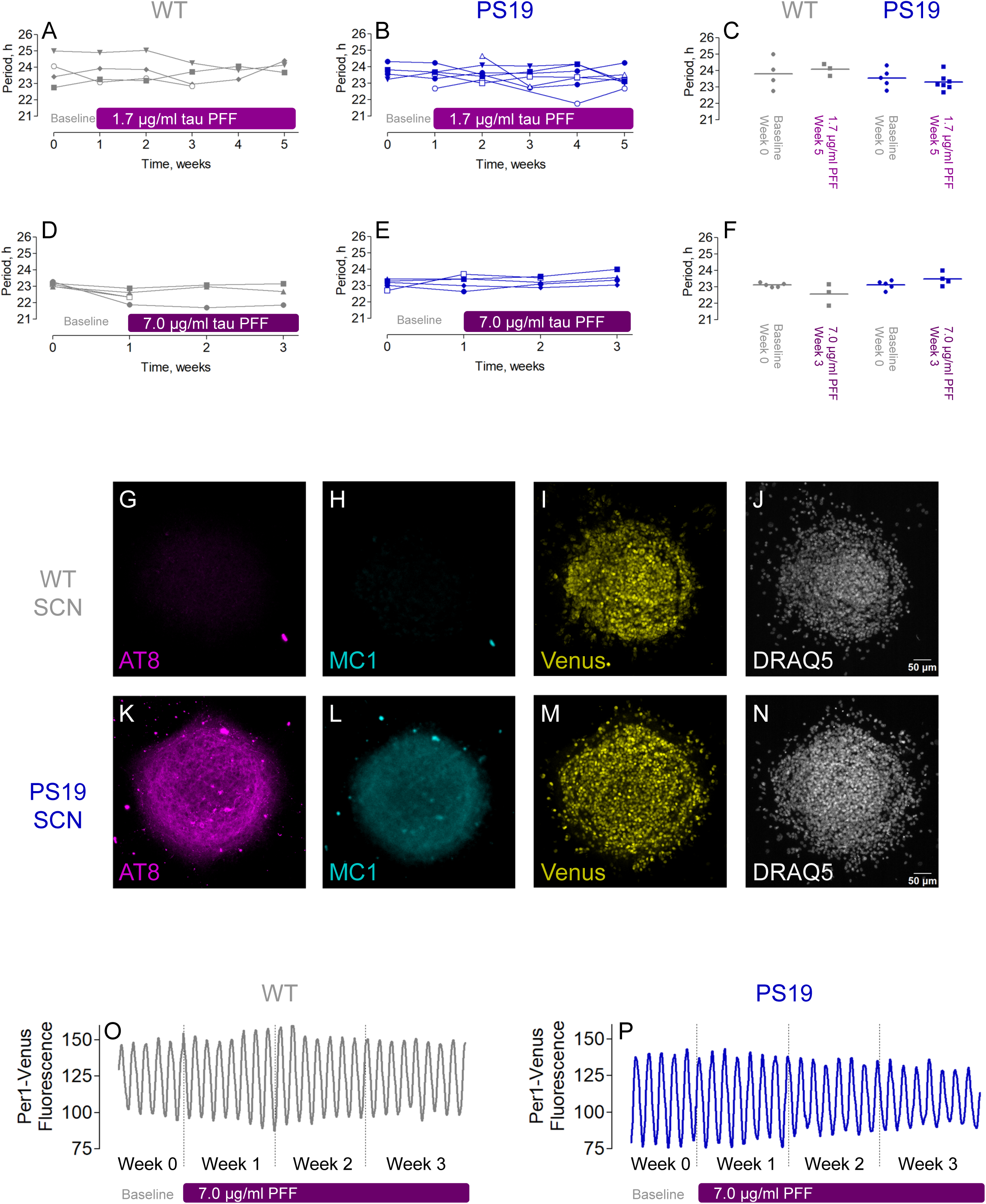
Profiling clock timing in SCN explant cultures. SCN explants were prepared from WT::*Per1*-Venus (WT) and PS19::*Per1*-Venus 1-day old pups and maintained in culture for several weeks on the incubator stage of a confocal microscope with Venus fluorescence acquisition occurring every hour. Baseline (Week 0) measurements were made on untreated cultures before the addition of either 1.7 µg/ml (A-C) or 7.0 µg/ml (D-F, O-P) tau PFF to the culture media. The circadian period was quantified in weekly intervals (from traces as depicted in O-P) and are plotted for individual SCN explants in A-B and D-E (open symbols indicate explants that have missing values either at the beginning or end of the acquisition). Scatterplots (C, F) of the baseline and final week period values for each tau PFF concentration depict a lack of statistically significant effect of treatment on circadian period (one-way ANOVA). Tau pathology in the PS19 explants treated with tau PFF (7.0 µg/ml) was confirmed by immunolabeling with AT8 (magenta pseudocolor) and MC1 (cyan pseudocolor) antibodies (G-H, K-L). Compared to WT, there was strong labeling with both pathologic tau antibodies, and individual cells with high levels of p-tau can be visualized. Venus immunolabeling and DRAQ5 (nuclear stain) labeling are included (I-J and M-N) for each genotype to demonstrate comparable levels of Venus expression and general explant morphology.

To accelerate tau pathology in the SCN explants and to further assess potential effects on cellular level clock timing, we extended these studies via the use of a tau seeding strategy, where tau PFF were added directly to the SCN explants. This approach followed the seeding strategy outlined by Guo and Lee (2013), where the addition of tau PFF led to a marked increase in fibril formation in cultured tissue from PS19 mice. After baseline data collection, PS19 and WT SCN explants were treated with 1.7 µg/ml tau PFF from Week 1 – Week 5 (Fig. 7A-C) or 7.0 µg/ml tau PFF from Week 1 – Week 3 (Fig. 7D-F and 7O-P) of Venus fluorescence acquisition. Following the seeding procedure, rhythms were profiled, and the period was quantified in weekly intervals. Despite robust levels of tau pathology markers (AT8 antibody labeling of p-tau and MC1 antibody labeling of pathological tau conformation) in PS19 SCN relative to WT (Fig. 7G-N), there were no differences in the periodicity of the rhythms across the two genotypes treated with tau PFF, again highlighting clock resilience despite pathological tau levels.

## 4. Discussion

Here, we tested for a functional or underlying relationship between tauopathy and the disruption of clock timing and entrainment properties of the SCN. Given the essential role of the SCN in circadian rhythm generation, coupled with prior work showing neurodegeneration and tau accumulation in the SCN of Alzheimer’s disease and frontotemporal dementia patients (Harper et al., 2008; Son et al., 2024a; Swaab et al., 1992), as well as work indicating SCN-related phenotypes in transgenic mouse models of tauopathy (Han et al., 2022; Stevanovic et al., 2017), we felt that a systematic examination of SCN clock timing and entrainment was merited. In the work reported here, a combination of *in vivo* and *in vitro* profiling studies did not detect a significant effect of tau accumulation or aggregation on the fidelity of the SCN clock.

Two possibilities may explain this lack of effect: 1) absent or insufficient transgene expression or 2) clock resilience to tau-pathology mediated perturbations.

The first possibility suggests that either the PS19 mouse model does not express the transgene in the SCN, or the expression level of pathologic tau is limited. However, our data and prior studies suggest that this is not the most compelling explanation.

In the PS19 mouse model, the *Prnp* promoter drives widespread neuronal expression of the human P301S tau transgene, resulting in pathological tau accumulation across multiple brain regions, including the hypothalamus (the location of the SCN; Chen et al., 2025; Ramirez et al., 2023; Sahara & Yanai, 2023; Wagner et al., 2015). Further highlighting the region’s susceptibility, the hypothalamus can accumulate tau pathology through propagation; injections of tau preformed fibrils (PFFs) into the hippocampus or striatum, induce the spread of tau pathology to the hypothalamus in PS19 mice (Iba et al., 2013).

It is also worth noting that hypothermia and weight loss are observed in PS19 mice (Patel et al., 2022). Given the critical role of the hypothalamus in regulating body temperature and metabolism, these data support the idea that key hypothalamus-regulated physiological processes are disrupted in the PS19 model. As such, these findings indicate that the PS19 mouse model expresses mutant human tau in the hypothalamus and that phenotypic effects on hypothalamic physiology can result.

Given these observations, it was somewhat unexpected to find that both clock timing and entrainment properties were not significantly affected in PS19 mice. Notably, our AT8 labeling revealed marked p-tau accumulation within the SCN as early as 2 months of age, preceding its appearance in several corticolimbic areas. Furthermore, our locomotor profiling studies assessed clock timing through the late stages of transgene-based neuropathology (10 months of age), when widespread CNS neurodegeneration and neuroinflammation are observed (Cao et al., 2023; Dutta et al., 2023; Guo et al., 2022; Huang et al., 2023; López-González et al., 2015; Patel et al., 2022; Wagner et al., 2015; Yoshiyama et al., 2010, 2007; Zhang et al., 2012). Nevertheless, SCN-based profiling did not reveal a significant phenotypic effect of mutant tau.

To further investigate this question, we used *in vitro* tau seeding methods developed by Guo and Lee (2013), in which we cultured SCN explants from PS19 animals, treated them with tau fibrils, and profiled for clock timing capacity. Even with clear evidence of cellular-level fibril formation (assessed using AT8 and MC-1 labeling), tissue-wide oscillatory capacity was not affected. Together, these data strongly suggest that even relatively high levels of tau accumulation do not disrupt SCN molecular clock timing properties.

Our findings are somewhat distinct from studies that reported alterations in the timing properties of the SCN (and/or clock gene expression) using several transgenic tauopathy models. Notably, Stevanovic et al. (2017) reported that 8-month-old Tg4510 mice exhibit greater locomotor activity during the day under a 12-h LD paradigm and longer free-running period than non-transgenic animals under DD conditions. In addition, rhythmic Period2 expression in the hypothalamus was not detected in Tg4510 mice. Further, in the P301S/PS19 model, altered phasing of the rhythm in BMAL1 in the SCN was reported, whereas peak Period2 expression was not affected (Han et al., 2022). These data indicate that tau pathology can alter aspects of clock timing and core clock gene expression within the SCN. With respect to the Stevanovic et al. (2017) study, potential differences in the reported phenotypic effects with the work reported here could have resulted from the use of different tau models (P301S/PS19 mouse line vs P301L/Tg4510 mouse line). In the Han et al. (2022) paper, which used PS19 mice, clock rhythms data were not reported. Here, it is also worth noting that mixed Aβ/tau models, such as the 3xTg line, have reported effects on clock timing at both the levels of the SCN and behavior (Sterniczuk et al., 2010; Weigel et al., 2023; Wu et al., 2018), whereas, in the TAPP model, alterations in clock-gated behavior were found to result from disrupted input to the SCN (Warfield et al., 2023). However, because Aβ and tau likely have complex additive or synergistic effects, it would be challenging to analyze our PS19 data within the context of studies that use bitransgenic Aβ/tau models.

The second possibility is that the SCN is resilient to perturbation, both at the level of core clock genes and the circuits that form the SCN oscillator. Regarding the molecular clock, redundancies within several components of the transcription-translation feedback loop (TTFL; including the *Period* and *Cryptochrome* gene families) allow the oscillator to function even when one homolog (e.g., *Period1* or *Period2*) is disrupted. In this regard, only *Bmal1* appears to be essential for timing; only its targeted deletion results in complete TTFL arrhythmia (Bae et al., 2001; Bunger et al., 2000; Cermakian et al., 2001; Horst et al., 1999; Pendergast et al., 2010).

Circuit-level resilience is also well-documented. Along these lines, selective lesioning studies found that overt rhythms can persist even when large portions of the SCN have been eliminated (Harrington et al., 1993; van den Pol and Powley, 1979). Further, work showing that the SCN has the capacity for functional rhythm recovery following partial SCN lesions highlights a remarkable degree of redundancy and plasticity within the SCN network (Eastman et al., 1984). Additionally, the SCN shows exceptional resistance to stress and excitotoxic cell death compared to other brain regions, such as the anterior hypothalamus and cortex. Studies using acute murine brain slices have shown that the SCN requires markedly higher NMDA concentrations to induce cell damage, even under conditions that cause significant injury in the anterior hypothalamus and cortex (Acharyya et al., 2022; Bottum et al., 2010). Interestingly, our immunolabeling data support this idea of SCN resilience. Hence, we did not detect gross anatomical changes in the SCN of PS19 mice, and both AVP- and VIP-expressing SCN neuronal populations were preserved (data were collected at 9 months of age when marked pathology has been reported throughout the CNS). Furthermore, the observation that both circadian rhythms and entrainment capacity were maintained in PS19 mice provides additional evidence that the SCN clock is highly functional in this model. However, it is important to note that postmortem analysis from human AD and FTD patients has detected neurodegeneration within the SCN (Harper et al., 2008; Son et al., 2024a; Stopa et al., 1999), and the loss of distinct cell populations was correlated with the reduced circadian drive (i.e., amplitude) over specific physiological processes. Clearly, with only one output measure used in the present study (locomotor activity), we might have missed potential dysregulation of other SCN-gated processes (e.g., temperature, hormone rhythms). Nevertheless, the data reported here showing intact and robust SCN clock rhythms and light entrainment raises important questions about the functional relationship between tauopathy and disruptions in the circadian timing system. A central question is centered on the neuroanatomical locus where clock-regulated physiological disruptions arise. The current results, showing intact SCN function despite pronounced tauopathy, suggest that the mechanistic link between tau pathology and circadian disruption may lie, in part, downstream of the SCN, within clock-gated systems and circuits (e.g., those regulating sleep and hormone rhythms).

Tau pathology appears to alter sleep/wake cycles and daily activity patterns-two processes that are influenced by circadian timing but are not direct outputs of the SCN itself (Anderson et al., 2009; Holth et al., 2017; Holton et al., 2020; Stevanovic et al., 2017). In line with this, our data support a model in which SCN function and output are preserved, but downstream clock-gated circuits governing behavior may be significantly disrupted by tauopathy, resulting in exacerbation of disease pathology. Further research using both animal models and human studies is needed to identify the precise mechanisms underlying this potential relationship. Finally, the possibility of a bidirectional interaction suggests that targeting circadian dysfunction could offer therapeutic benefits in managing tauopathies. Understanding this connection is critical, as circadian disruption may not only be a symptom of tauopathy but could also contribute to its progression.

---

**Supplemental Table 1.** Analysis of circadian locomotor activity parameters by sex. All parameters indicated in the “Intermediate-Late” stage pathology section (see Table 1) were analyzed for sex-related differences. “Sex p-values” were calculated by comparing sexes within the same genotype (e.g., WT male vs WT female). “Genotype p-values” were calculated by comparing the same sex between genotypes (e.g., WT male vs PS19 male). Data are presented as the mean ± SEM for all values. * = p < 0.05, Student’s t-test. Statistically significant sex-specific differences are noted for WT animals (parameters 7 & 8); No statistically significant sex-specific differences in circadian parameters are noted for PS19 mice.

| Parameter<br>(mean ±<br>standard<br>error) | WT male (n) | WT Female (n) | PS19 male (n) | PS19 female (n) | Sex *P*-Value<br>(M v F) | Genotype *P*-Value<br>(WT v PS19) |
| --- | --- | --- | --- | --- | --- | --- |
| 1. Overall activity in LD<br>(counts/5 min) | 78.57 ± 12.75 (8) | 80.90 ± 20.79 (6) | 83.97 ± 13.37 (8) | 89.41 ± 22.67 (6) | **WT:** 0.922<br>**PS19:** 0.831 | **Male:** 0.774<br>**Female:** 0.788 |
| 2. LD Phase Angle of activity onset relative to lights off (min) | -2.05 ± 7.64 (8) | -11.77 ± 7.39 (6) | -12.13 ± 9.36 (8) | -10.37 ± 8.52 (6) | **WT:** 0.391<br>**PS19:** 0.896 | **Male:** 0.418<br>**Female:** 0.904 |
| *Negative values indicate activity onset prior to “lights off”* |  |  |  |  |  |  |
| 3. Days to re-entrain to 6h advancing LD cycle (d) | 5.00 ± 0.87 (7) | 6.00 ± 0.71 (4) | 4.4 ± 1.03 (5) | 4.20 ± 0.73 (5) | **WT:** 0.457<br>**PS19:** 0.878 | **Male:** 0.666<br>**Female:** 0.127 |
| 4. Days to re-entrain to 6h delaying LD cycle (d) | 2.63 ± 0.57 (8) | 4.20 ± 1.28 (5) | 3.17 ± 0.83 (6) | 2.33 ± 0.71 (6) | **WT:** 0.225<br>**PS19:** 0.465 | **Male:** 0.587<br>**Female:** 0.215 |
| 5. Overall activity in DD<br>(counts/5 min) | 72.09 ± 10.33 (7) | 72.86 ± 13.74 (6) | 70.06 ± 15.76 (6) | 87.93 ± 18.59 (6) | **WT:** 0.964<br>**PS19:** 0.480 | **Male:** 0.914<br>**Female:** 0.529 |
| 6. Circadian period (τ) during DD | 23.55 ± 0.06 (7) | 23.63 ± 0.06 (6) | 23.51 ± 0.08 (6) | 23.56 ± 0.08 (6) | **WT:** 0.339<br>**PS19:** 0.699 | **Male:** 0.734<br>**Female:** 0.483 |
| 7. Duration of alpha during DD (h) | 7.82 ± 0.46 (7) | 9.41 ± 0.52 (6) | 8.54 ± 0.97 (6) | 8.82 ± 0.63 (6) | **WT:** 0.043*<br>**PS19:** 0.815 | **Male:** 0.497<br>**Female:** 0.484 |
| 8. Acrophase during DD (CT) | 16.21 ± 0.19 (7) | 17.11 ± 0.19 (6) | 16.69 ± 0.39 (6) | 16.68 ± 0.34 (6) | **WT:** 0.007*<br>**PS19:** 0.990 | **Male:** 0.265<br>**Female:** 0.305 |

## Notes

**Conflict-of-interest statement**: The authors declare that they have no conflicts of interest related to this work.

Funding Sources: National Institutes of Health: GM133032, AG065830

### Competing Interest Statement

The authors have declared no competing interest.

